# Optimizing the Martini 3 force field reveals the effects of the intricate balance between protein-water interaction strength and salt concentration on biomolecular condensate formation

**DOI:** 10.1101/2022.11.07.515502

**Authors:** Gül H. Zerze

## Abstract

Condensation/dissolution has become a widely acknowledged biological macromolecular assembly phenomenon in subcellular compartmentalization. MARTINI force field offers a coarse-grained protein model with a resolution that preserves molecular details with an explicit (CG) solvent. Despite its relatively higher resolution, it can still achieve condensate formation in reasonable computing time with explicit solvent and ionic species. Therefore, it is highly desirable to tune this force field to be able to reproduce the experimentally observed properties of the condensate formation. In this work, we studied the condensate formation of the low-sequence complexity (LC) domain of FUsed in Sarcoma (FUS) protein using a MARTINI 3 force field by systematically modifying (increasing) the protein-water interaction strength and varying salt concentration. We found that the condensate formation is sensitive both to the protein-water interaction strength and the presence of salt. While the unmodified MARTINI force field yields a complete collapse of proteins into one dense phase (i.e., no dilute phase), we reported a range of modified protein-water interaction strength that is capable of capturing the experimentally found transfer free energy between dense and dilute phases. We also found that the condensates lose their spherical shape upon the addition of salt, especially when the protein-water interactions are weak. Inter-chain amino acid contact map analysis showed one explanation for this observation: the protein-protein contact fraction reduces as salt is added to systems (when the protein-water interactions are weak), consistent with electrostatic screening effects. This reduction might be responsible for the condensates becoming nonspherical upon the addition of salt by reducing the need for minimizing interfacial area. However, as the protein-water interactions become stronger to the extent that makes the transfer free energy agree well with experimentally observed transfer free energy, we found an increase in protein-protein contact fraction upon the addition of salt, consistent with the salting-out effects. Therefore, we concluded that there is an intricate balance between screening effects and salting-out effects upon the addition of salt and this balance is highly sensitive to the strength of protein-water interactions.

## Introduction

The condensation/dissolution phenomenon in subcellular compartmentalization has been brought to the spotlight by the seminal work by Brangwynne et al. for cytoplasmic granules,^1^ after which the research field of biomolecular organization has exploded with many other examples of subcellular compartments that form via a similar phenomenon.^2–4^ One mechanism that explains this phenomenon is liquid-like phase separation (LLPS),^5^ in which the two liquid-like phases are a protein-rich (condensed or dense) phase and a dilute phase. Phase transitions, like LLPS, are ubiquitous in biological macromolecular assembly, and vital for organizing and regulating cellular biochemical processes.^6–13^

Molecular simulations of biomolecular condensates have been particularly useful in providing means to directly probe the interactions that mediate the condensate formation. The fundamental trade-off in molecular modeling (i.e., the accessible time and length scales versus the model resolution), however, limits the range of applications of a given model, e.g., resolutions lower than atomistic scale are often needed for systematic studies of condensate formation.^14^ Accordingly, coarse-grained (CG) models have become a popular option to study condensate formation.^15–25^ Among the CG models, the MARTINI force field offers a resolution that preserves molecular details with an explicit (CG) solvent while being able to achieve condensate formation in reasonable computing time.^22,24^

The MARTINI force field has originally been developed to reproduce experimental free energies of chemical building blocks’ partitioning between polar and apolar phases^26,27^ and has been employed successfully for several purposes. Some recent examples include lipid nanoemulsions for drug delivery,^28^ PEGylation of proteins,^29^ predicting the structure of membrane-associated protein assemblies.^30^ Recently, Benayad et al.^24^ has shown that an older version of the MARTINI force field (version 2.2) can also successfully reproduce the experimentally observed (*in vitro*) excess transfer energy of the low-sequence complexity domain of FUS protein (FUS LC) between condensed and dilute phases by scaling the proteinprotein interactions. The authors obtained further insights into the condensate properties by using this scaled MARTINI 2.2 force field, such as shear viscosity and surface tension of droplets. Moreover, Tsanai et al.^22^ have shown that an open beta version of MARTINI 3.0 can reproduce the salt-dependent coacervate formation of polyelectrolyte type peptides (by applying elastic bonds between backbone beads to enforce backbone extension). In this work, we aim to optimize an open-beta version of the MARTINI 3.0 force field to reproduce the experimentally found properties of the biomolecular condensate formation.

Single-molecule properties, most notably, the size-related measurables (such as radius of gyration) have been successfully used as objective functions for force field optimization, which yield a reasonable agreement between experiments and simulation data for non-size-related test parameters (such as backbone structure, transport properties, etc.)^31–34^ One of the most commonly used tuning parameters for such force field optimizations is the protein-water interaction strength (specifically, *ϵ* of Lennard-Jones [LJ] interactions between protein-water atoms/particles). This parameter has been successfully used for the optimization of atomistic force fields^31,32^ as well as the MARTINI force field.^33,35^ Larsen et al.^33^ and Thomasen et al.^35^ have devised this approach for an open-beta version of MARTINI 3.0 (3.0.beta.4.17) and MARTINI 3.0.0 (release version), respectively, where they showed that overly compact behavior of intrinsically disordered proteins and multidomain proteins can be improved by 6 to 10% increase in protein-water interactions.

In this work, we have employed a similar approach as in the work by Lindorff-Larsen^33,35^ by systematically modifying the nonbonded (LJ) interaction strength between the protein-water beads, in order to reproduce the condensate properties, specifically the excess free energy of transferring protein chains from dilute phase to condensed phase.^24^ For this benchmarking study, we used a 163 residue-long peptide (FUS LC) that represents the low sequence-complexity domain of the FUS RNA-binding protein. We presented the results from each scaling at various salt concentrations. We found that the dense phase formation is sensitive to both the strength of protein-water interaction and the salt concentration. We reported a range of protein-water interaction scaling where we were able to reproduce experimentally observed transfer free energy although we found that the protein concentration in the dense phase was higher than the experimentally detected concentrations. We also found that the presence of salt concentration affects the dense phase morphology, especially when the protein-protein interactions dominate the protein-water interactions. We showed that the individual amino acid contacts are highly sensitive to salt concentration; salt has more predominant screening effects when protein-water interactions are weak whereas the opposite salting-out effects become more predominant as protein-water interactions strengthen. We conclude that molecular simulations with well-balanced force fields (i.e., well-balanced protein-protein and protein-water interactions) are highly useful in studying the microscopic details of the interactions governing biomolecular self-assembly.

## Methods

Here we performed coarse-grained molecular dynamics simulations of the FUS LC domain which is an ideal benchmarking system as it has been shown to form condensates *in vitro* ^36^ and has plenty of experimental and computational data available for comparison.^36–39^ The full sequence of simulated 163 residue-long FUS LC is given in Supporting Information (SI).

### Modeling

We employed an open beta version of the MARTINI 3.0 force field (3.0.beta.3.2)^40,41^ for modeling the FUS LC domain. We kept the protein-protein interactions unmodified (and no additional elastic backbone constraints were applied) but tested a range of scaled protein water interactions. Specifically, we modified the ε of the Lennard-Jones interactions between protein and water beads following a strategy similar to the ones in the single-molecule studies by Lindorff-Larsen and co-workers.^23,33,35^ We will refer to the scaling parameter as *λ*; the parameter with which we multiply the epsilon of Lennard-Jones interactions between protein and water beads. We presented the results where the *λ* parameter varies between 1-1.04.

Initial atomic coordinates of FUS LC were prepared as an extended configuration that is energy minimized using Amber99SBdisp force field.^42^ The atomistic coordinates were then converted to coarse-grained coordinates by using the *martinize* script provided within MARTINI 3.0.beta.3.2 distribution.^41^ We prepared cubic simulation boxes (L = 40 nm) that contained 100 copies of FUS LC chains. This setting gives a protein concentration of ≈2.6 mM (44.5 mg/mL). Benayad et al. have shown that this concentration fits within the range that would yield a spherical condensate (instead of a percolated network).^24^ Moreover, for this volume and concentration, spherical droplet formation simulations and direct coexistence (slab) simulations yield very similar equilibrium densities of the condensed and dilute phases.^24^

Each of our systems starts from a monodispersed initial condition. A snapshot of an example starting configuration is shown in the SI (Figure S1). The simulation boxes were then solvated in MARTINI water and randomly selected water beads are replaced with ion beads both to provide electroneutrality (a single chain of FUS LC domain has a net negative charge −2) and to adjust the salt concentration (tested 0, 50, 100, and 200 mM NaCl concentrations). We used the “tiny” bead type (TQ1) both for Na+ and Cl- ions and kept the ions-water and ion-protein interactions unmodified.

### Simulation Details

After initial solvation, all systems were energy minimized using the steepest descent algorithm. They were then equilibrated with 400 ps NVT simulations (T=300 K) followed by 400 ps NPT simulations (T = 300 K, P = 1 bar), where the temperature was maintained constant with the velocity-rescaling algorithm^43^ a 2 ps time constant and pressure was maintained constant using Parrinello-Rahman barostat^44^ with a time constant of 20 ps. Electrostatic interactions were calculated using a generalized reaction field method.^45^ A cutoff distance of 1.1 nm was used both for the van der Waals interactions and electrostatic interactions. The leapfrog algorithm was used to integrate the equations of motion with a time step of 20 fs. All simulations were performed using GROMACS MD engine (version 2016.3).^? ?^

### Cluster Formation Analysis

Cluster formation is defined based on a minimum distance criterion. We first calculated the closest distance between each protein pair as a function of time. Any two protein molecules are considered to be in the same cluster if any two beads of the molecules are within 0.5 nm (or less) distance from each other. Based on this criterion, we built adjacency matrices and then found the connected components by using the compressed sparse graph routines of public Python libraries.^46^ We then labeled the connected components based on their size; the largest one being the largest cluster and so on.

## Results and Discussions

### Identification of condensate formation

We first identified the dense-phase (droplet) formation by analyzing the cluster formation during the simulations. Any two protein molecules are considered to be in the same cluster if any two beads of the molecules are within 0.5 nm (or less) distance from each other (see Methods for details). We then quantified the fraction of proteins within given clusters. Fractions of proteins in the most populated three clusters are presented in Figure 1. While all the systems start from a randomly monodispersed state, we found that all protein chains coalesce into a single cluster for smaller *λ* (*λ* < 1.02) values after some equilibration time (Figure 1). We determined the equilibration time based on the cluster formation (Figure 1) and the fluctuations in the total potential energy of the system (Figure S2). Equilibration periods are tabulated in Table S1 for all systems. Representative snapshots of the systems are shown in Figure 2 after the cluster formation was observed (i.e. after the equilibration time).

**Figure 1:**
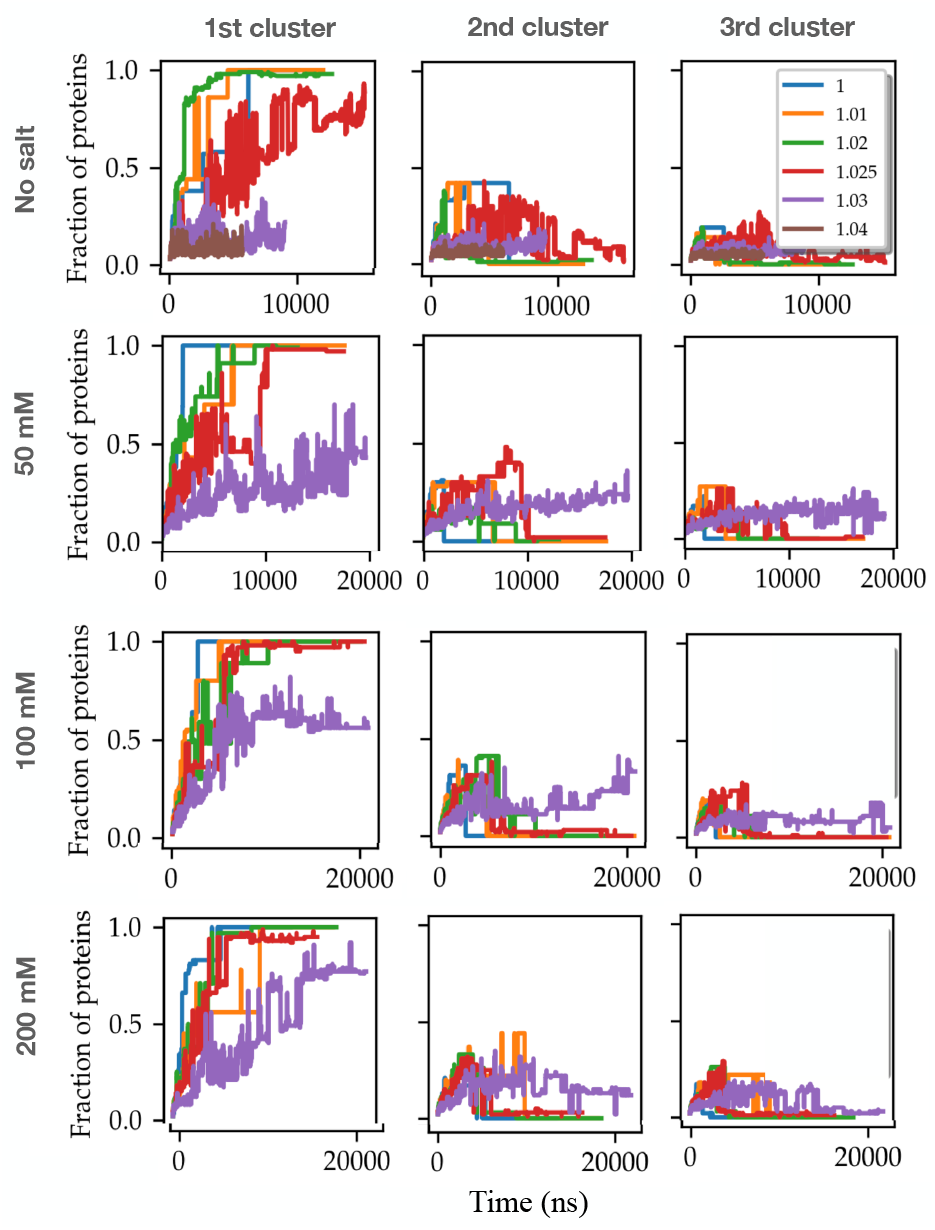
Fraction of proteins in three most populating clusters for the *λ* values tested and for various salt concentrations (top row: 0 mM salt, second row: 50 mM NaCl, third row 100 mM NaCl, last raw: 200 mM NaCl)

**Figure 2:**
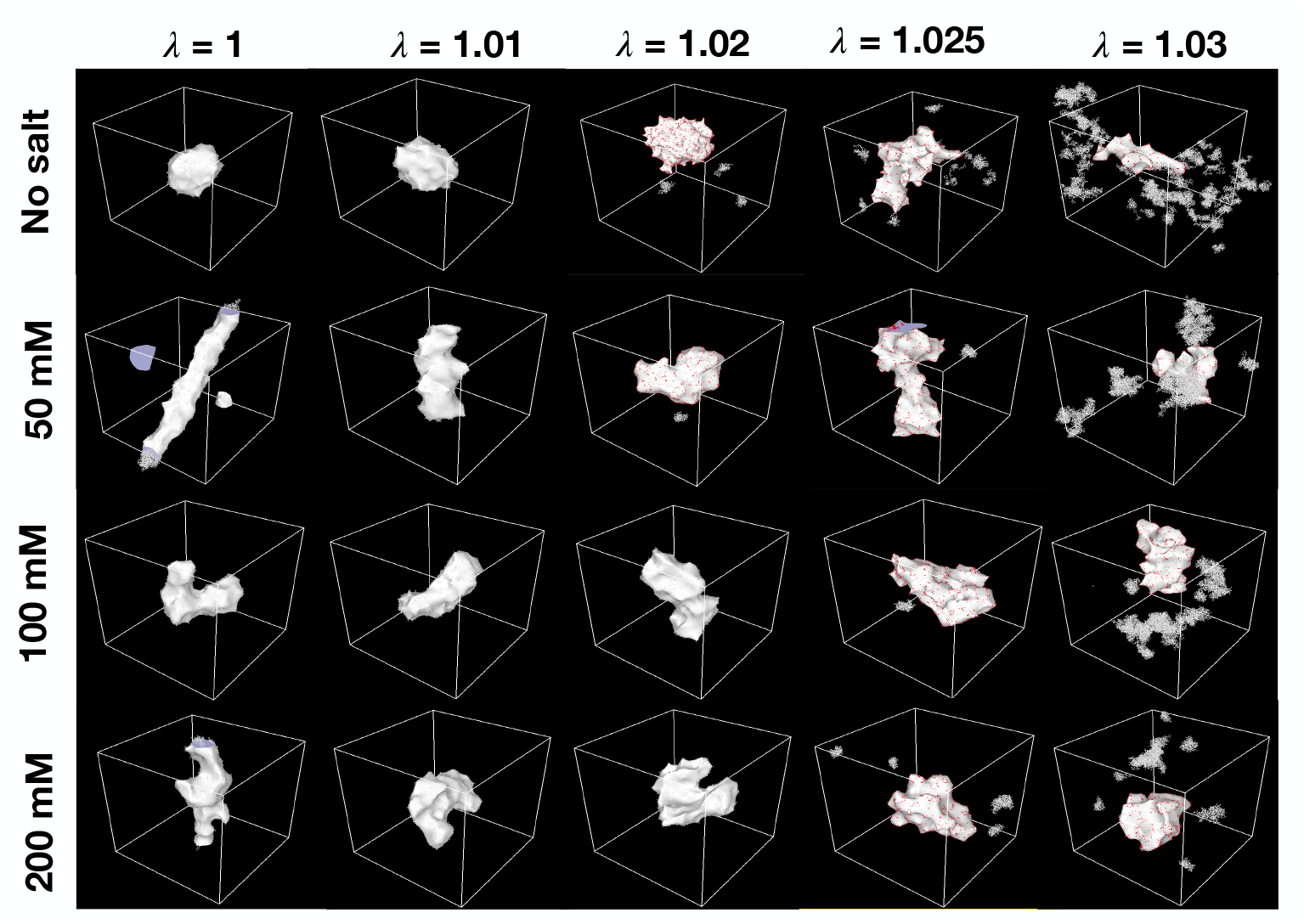
Representative snapshots of the systems for the *λ* values tested and for various salt concentrations (top row: 0 mM salt, second row: 50 mM NaCl, third row 100 mM NaCl, last raw: 200 mM NaCl) obtained after the cluster formation is observed. A surface representation (with a probe radius 8Å) is used to illustrate protein in OVITO. Further details of the surface reconstruction are available in *Concentration profiles* subsection.

We labeled the cluster with the largest fraction of proteins as the droplet (or the dense phase) and the remainder of the system as the dilute phase. We did not find any proteins in the dilute phase for *λ* < 1.02. At no salt condition, for *λ* = 1.02, we found that while the majority of the proteins (97-98%) coalesce into one cluster, a small percentage of proteins (fluctuating between 2 to 3%) is present in the dilute phase (Figure 1). The fraction of proteins in the largest cluster (i.e., in the dense phase) becomes smaller as *λ* gets larger. At *λ* = 1.025 (no salt), this fraction becomes ≈ 0.75 and the droplets start to show significant deviations from a round shape (see the snapshot given in Figure 2, top row, the fourth column from left). Beyond *λ* = 1.03 there was not a dominant cluster formation (Figure 1).

For the conditions where we have a finite salt concentration, we found that clusters are forming beyond the periodic boundaries (percolation) for some cases, especially at small *λ* (e.g., see Figure 2, second row, the first column from left for *λ* = 1 at 50 mM salt concentration), unlike no salt conditions. These clusters at finite salt concentrations also show substantial deviations from a spherical shape. These observations indicate that the condensate morphologies change significantly upon the addition of salt. While these changes in the morphology are a subject of future work, here we further analyzed the per-residue contact formation between the protein chains in the *Contact formation* subsection, which could shed some light on the impacted morphologies.

### Concentration profiles

In order to analyze the partitioning of individual species into the condensed and dilute phases, we quantified the average radial concentration profiles of the species. For this analysis, we defined the center as the center of the condensate; calculated the concentration profiles for each instantaneous frame (after equilibration); and calculated the ensemble average. While the protein concentration profile for no salt conditions follows a sigmoidal trend, which is consistent with their sphere-like shape, that for finite salt concentrations do not follow a clear sigmoidal trend. The lack of a sigmoidal trend is also consistent with their condensate’s substantial deviations from a spherical shape (Figure 2). For no salt conditions, we fit the data to a sigmoidal function as given in Eq. 1.

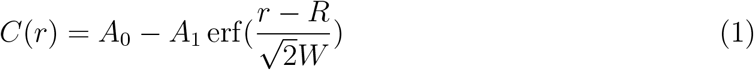

r is the radial distance from the droplet center; erf is the error function; and *A*_0_, *A*_1_, *R*, and *W* are fit parameters. As noted in the work by Hummer and coworkers,^24^ the density plateaus at the center for sufficiently large droplets with a defined density, where *A*_0_ + *A*_1_ and *A*_0_ – *A*_1_ will be the limiting concentrations, dense and dilute phase concentrations, respectively. For no salt conditions, we were able to estimate the dilute and condensed phase concentrations from these limiting values. However, since the condensates for the conditions with finite salt concentrations significantly deviate from a sphere (and they do not show a clear plateau as the center is approached), we used a surface reconstruction method to estimate the volume and concentration as an alternative approach.^47^ In this approach, a virtual probe is used to define the accessible spatial volume, the inaccessible volume, and the surface between the two. We used OVITO software for this analysis.^48^ We chose a 10Å radius for the probe, which takes the nearest bead separation within the condensed phase into account. We also have tested a range of radii between 8Å to 16Å for no salt conditions and found that the difference between the condensed phase concentration estimations from two different methods is smallest for 10Å probe for the small *λ* (Figure S3). In Table 1, condensed phase concentrations (obtained from two different techniques for no salt) are tabulated.

**Table 1:**
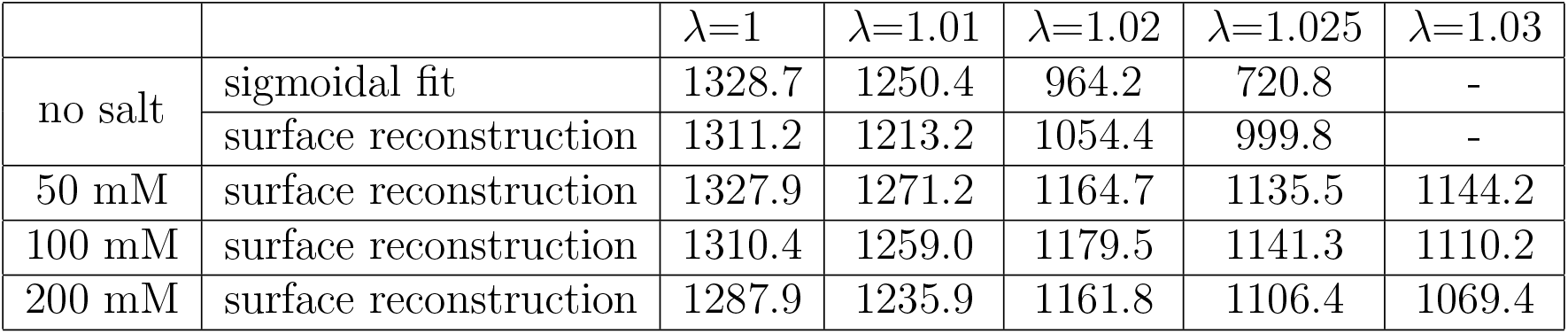
Protein concentrations in the dense phase are given in mg/mL. For the no-salt condition, the concentrations are obtained from two different techniques (sigmoidal fit and surface reconstruction, see the text for more details). Statistical errors calculated by block averaging of the data (dividing the equilibrated data into two equal blocks) are less than 1%.

| | | $\lambda=1$ | $\lambda=1.01$ | $\lambda=1.02$ | $\lambda=1.025$ | $\lambda=1.03$ |
| --- | --- | --- | --- | --- | --- | --- |
| no salt | sigmoidal fit | 1328.7 | 1250.4 | 964.2 | 720.8 | - |
|  | surface reconstruction | 1311.2 | 1213.2 | 1054.4 | 999.8 | - |
| 50 mM | surface reconstruction | 1327.9 | 1271.2 | 1164.7 | 1135.5 | 1144.2 |
| 100 mM | surface reconstruction | 1310.4 | 1259.0 | 1179.5 | 1141.3 | 1110.2 |
| 200 mM | surface reconstruction | 1287.9 | 1235.9 | 1161.8 | 1106.4 | 1069.4 |

We report the protein concentration in the dense phase only until *λ* = 1.03 since we do not observe a viable cluster formation beyond *λ* = 1.03. As *λ* increases, the volume of the dense phase increases (and condensed phase concentration decreases accordingly) until the system is not capable of forming a dense phase (*λ* > 1.03). The smallest protein concentration that we obtain in the condensed phase is 999.8 mg/mL (using surface reconstruction; or 720.8 using sigmoidal fit). We note that regardless of the analysis technique we used, even the smallest protein concentration that we obtain is still larger than the experimentally found protein concentration in the condensed phase. We discuss this further in the following subsection. We also report the concentrations in the dilute phase in Table 2.

**Table 2:** Protein concentrations in the dilute phase given in mg/mL (surface reconstruction).

| | $\lambda=1$ | $\lambda=1.01$ | $\lambda=1.02$ | $\lambda=1.025$ | $\lambda=1.03$ |
| --- | --- | --- | --- | --- | --- |
| no salt | 0 | 0 | 1.4 | 10.0 | - |
| 50 mM salt | 0 | 0 | 1.7 | 11.3 | 30.8 |
| 100 mM salt | 0 | 0 | 3.6 | 1.3 | 17.3 |
| 200 mM salt | 0 | 0 | 0 | 2.7 | 10.7 |

As can be seen in Table 2, there is no protein in the dilute phase for *λ*=1 and *λ*=1.01 (for any salt condition), i.e., all proteins are in the dense phase. Coexistence of dense and dilute phases applies for *λ*=1.02 and 1.025 for no salt conditions. Beyond *λ*=1.03 there is no dense phase formation (for no salt conditions, there is no dense phase formation for *λ*=1.03). The relative concentrations in the dense and dilute phase are discussed further in the following subsection.

We also found that water fraction (mass) in the dense phase increases as *λ* increases (Figure 3B). We found the largest water fraction For the ionic species, we found that while they are concentrated in the dense phase (Figure 3C), there still is some ionic concentration in the dilute phase (including *λ*=1). In summary, protein-protein interactions dominate the protein-water interactions at smaller *λ* values, yielding very high protein concentrations in the dense phase and very small to no protein concentrations in the dilute phase. Since our goal is to optimize the *λ* until we establish a balance between protein-protein and protein-water interactions, we next calculated the excess transfer free energy based on the relative concentrations and compared our findings against experimental data.

**Figure 3:**
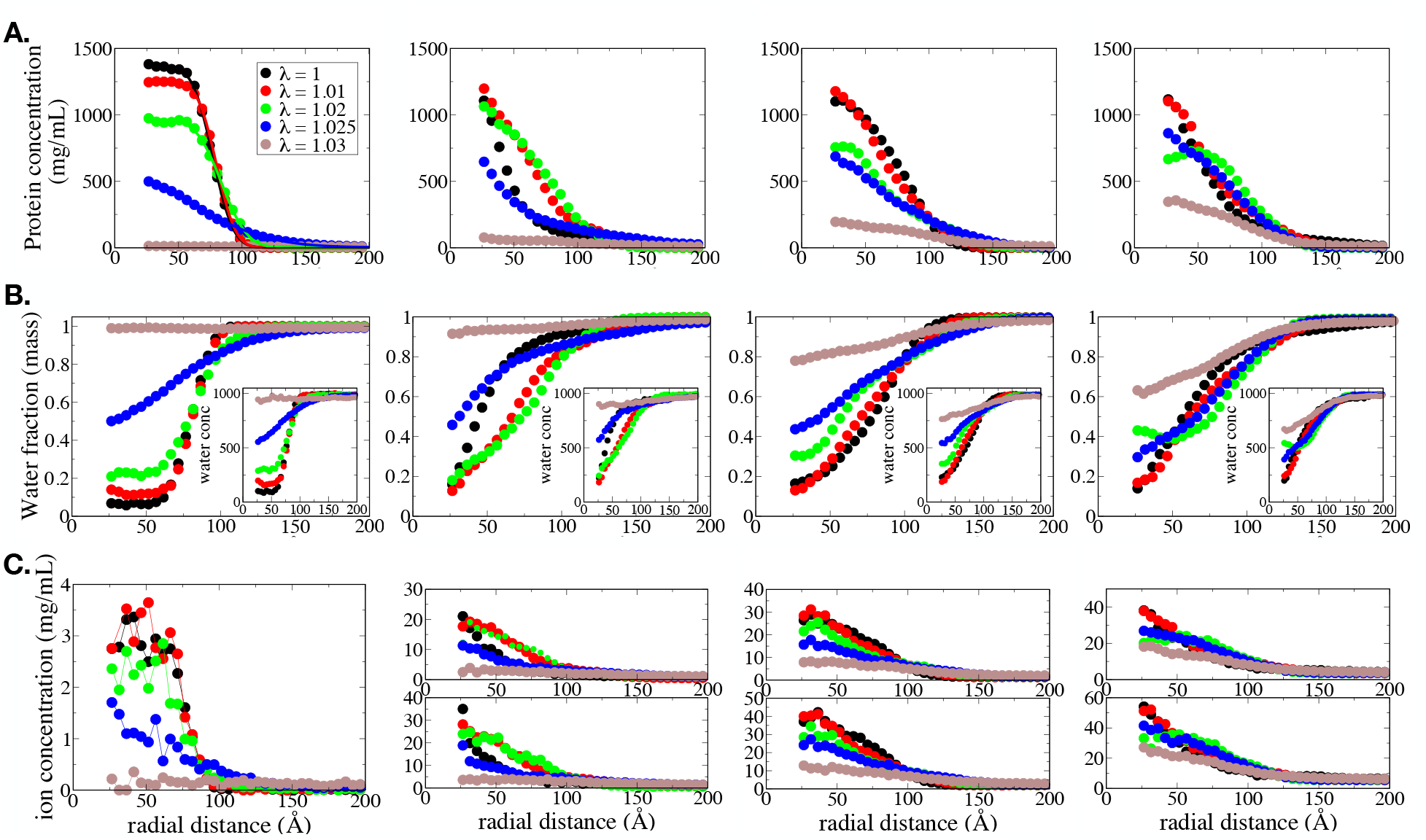
Radial concentration profiles of protein (A), water (B), and ions (C). From left to right, first, second, third, and fourth columns are no salt, 50 mM salt, 100 mM salt, and 200 mM salt concencentrations, respectively.

### Excess transfer free energy and relative concentrations of the condensed and dilute phases of FUS LC condensates

Phase diagrams are quantitative descriptors of LLPS by defining the thermodynamic conditions at which distinct phases form (e.g. condensed and dilute phases) and coexist at equilibrium. Phase diagrams of LLPS are often defined on the temperature-concentration plane. Accordingly, coexistence data based on relative concentrations of condensed and dilute phases at a given temperature and pressure are metrics of particular interest. Benayad et al.^24^ used a metric that takes this relative concentration into account to benchmark the scaled protein-protein interactions in MARTINI 2 force field. They have used excess transfer energy, that is, the free energy associated with the transfer of protein chains from the dilute phase to the condensed droplet phase, formulated as:

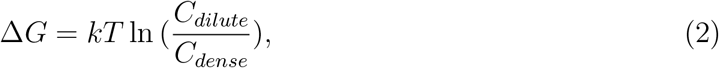

where *k* is the Boltzmann constant, *T* is the temperature, and *C_dilute_* and *C_dense_* are the concentrations in the dilute and dense phases, respectively. In their work, Benayad et al.,^24^ for atmospheric pressure and room temperature, have identified a range of protein-protein scaling parameters which yield coexistence concentrations that agree well with the experimental data. In this work, we reported a similar experimental comparison for the scaled protein-water interactions. In Figure 4A, we presented the protein concentration in the dense phase for different salt concentrations and scaling parameters (*λ*) that we tested. As noted in the previous section, dense phase volumes are estimated from a surface reconstruction technique. The general trend is a decrease in the dense phase protein concentration as *λ* increases, except for *λ*=1.03 (smallest *λ* that forms a dense phase). Although our dense phase concentrations are larger than the experimentally identified dense phase concentrations (Figure 4A), we still found a range of protein-water scaling parameters for which the excess transfer free energy agrees well with the experimental data (Figure 4B) within the range that we observe phase coexistence (1.01 < *λ* ≤ 1.03). The experimental data is compiled from the work by Fawzi and coworkers.^36,39^

**Figure 4:**
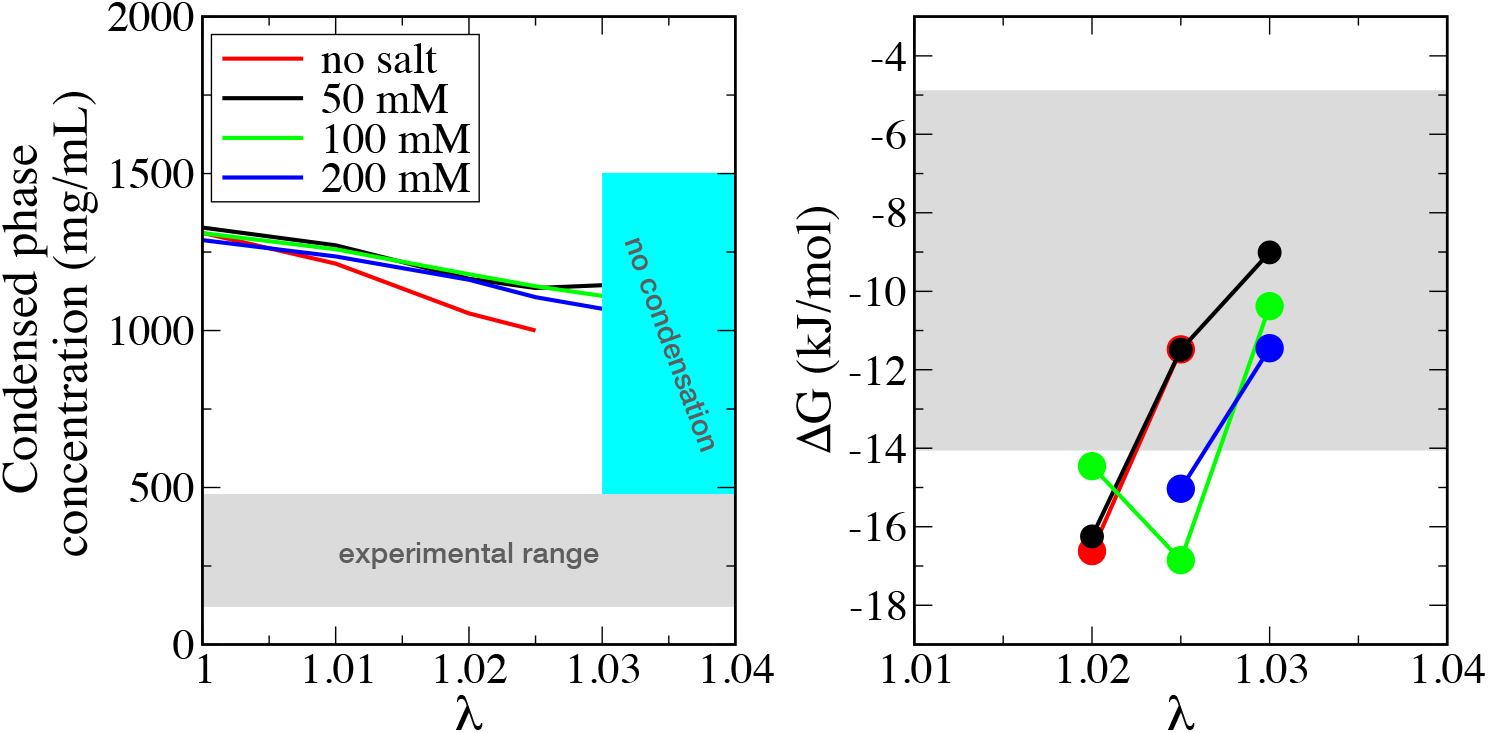
Protein concentrations in the dense phase as a function of the scaling parameters (*λ*). Gray-shaded areas are experimentally found ranges. See the text for experimental reference.

### Contact maps

We also calculated the inter-chain residual contact maps (square, symmetric matrices) at each *λ* parameter and salt concentration to quantitatively visualize individual contacts between different amino acids and contact clusters that hold the condensates together. Figure 5 reports the ensemble-averaged fraction of contacts between each residue *i, j*. We will analyze the data in Figure 5 in two dimensions: i) as a function of the *λ* parameter, and ii) as a function of salt concentration. Figure 5 shows that the contact propensity decreases as the *λ* parameter increases (from left to right), that is, as the protein-water interaction strengthens, at all salt concentrations. This observation is consistent with our findings reported in previous sections that the cluster formation weakens as the *λ* parameter increases at all salt concentrations. As the protein-protein contact formation weakens, condensate formation also weakens. The majority of the strong contacts (fraction > 0.1) are the tyrosine residues in (S/G)Y(S/G) repeating units of the FUS LC domain (Figure 6B). The 163 residue-long LC domain has 18 of these repeating units (SI Text). It has been previously shown that a progressive mutation of the central tyrosine residues to serine in (S/G)Y(S/G) consensus motif causes a systematic drop in the partitioning of the FUS LC domain to the stress granules.^37^ Our findings are consistent with this experimental observation. We found that the lower the *λ* is (i.e., the larger the fraction of proteins in the largest cluster), the stronger the interactions of the central tyrosine residues in the (S/G)Y(S/G) consensus motif. Accordingly, we conclude that these central tyrosine residues play a major role in driving the dense phase formation.

**Figure 5:**
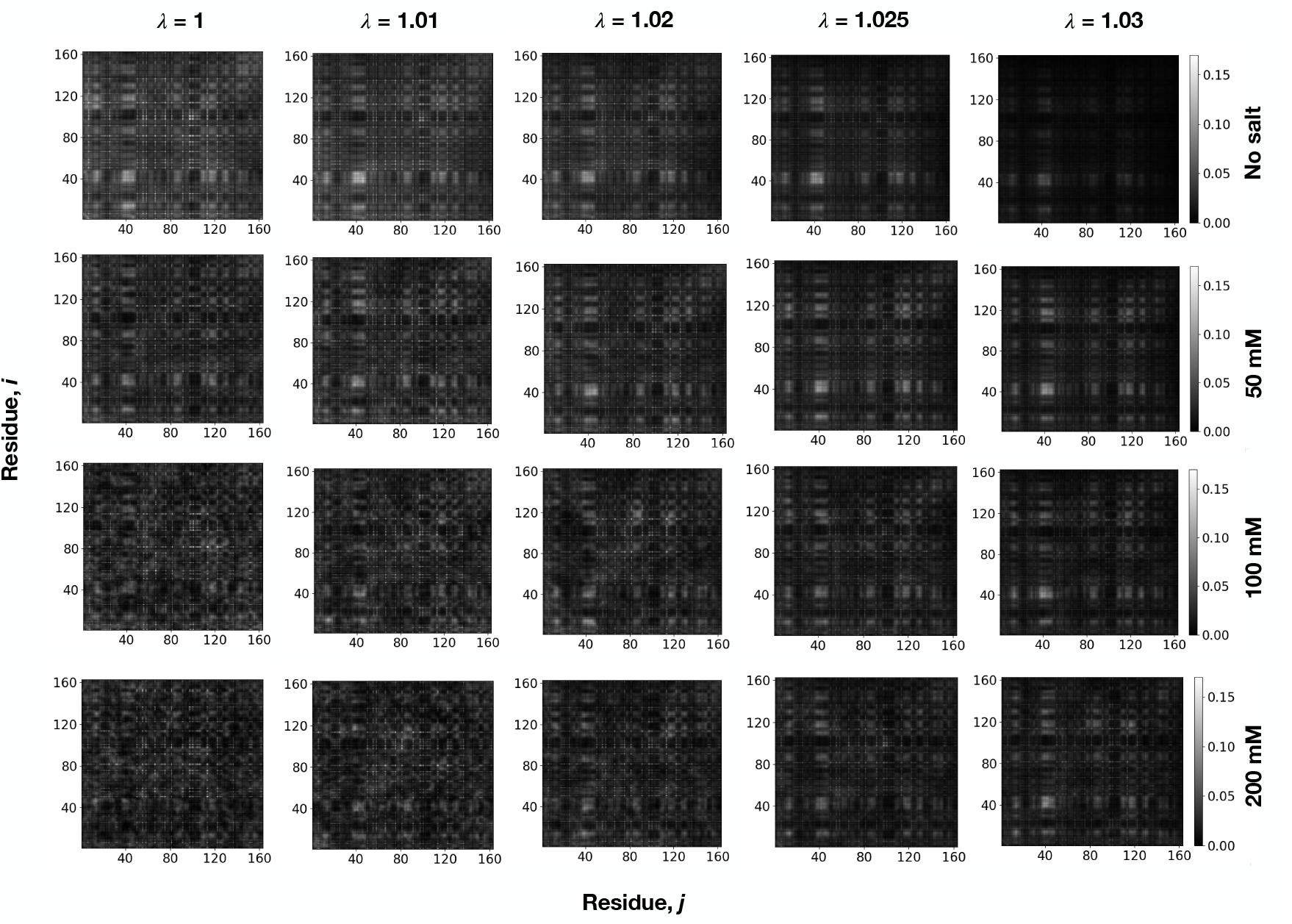
Fraction of the inter-chain residual contacts between each residue *i, j*. We first calculated a 16300×16300 binary matrix of contacts at each instantaneous frame (using a 0.8 nm cutoff), considering all protein chains in the system (we have 100 identical chains, each one is 163 residues). For each frame, we first averaged the contacts over 100 chains, reducing the matrix size to 163×163, and then finally reported the ensemble average as the *fraction* of contact formation (the color scale from black to white).

**Figure 6:**
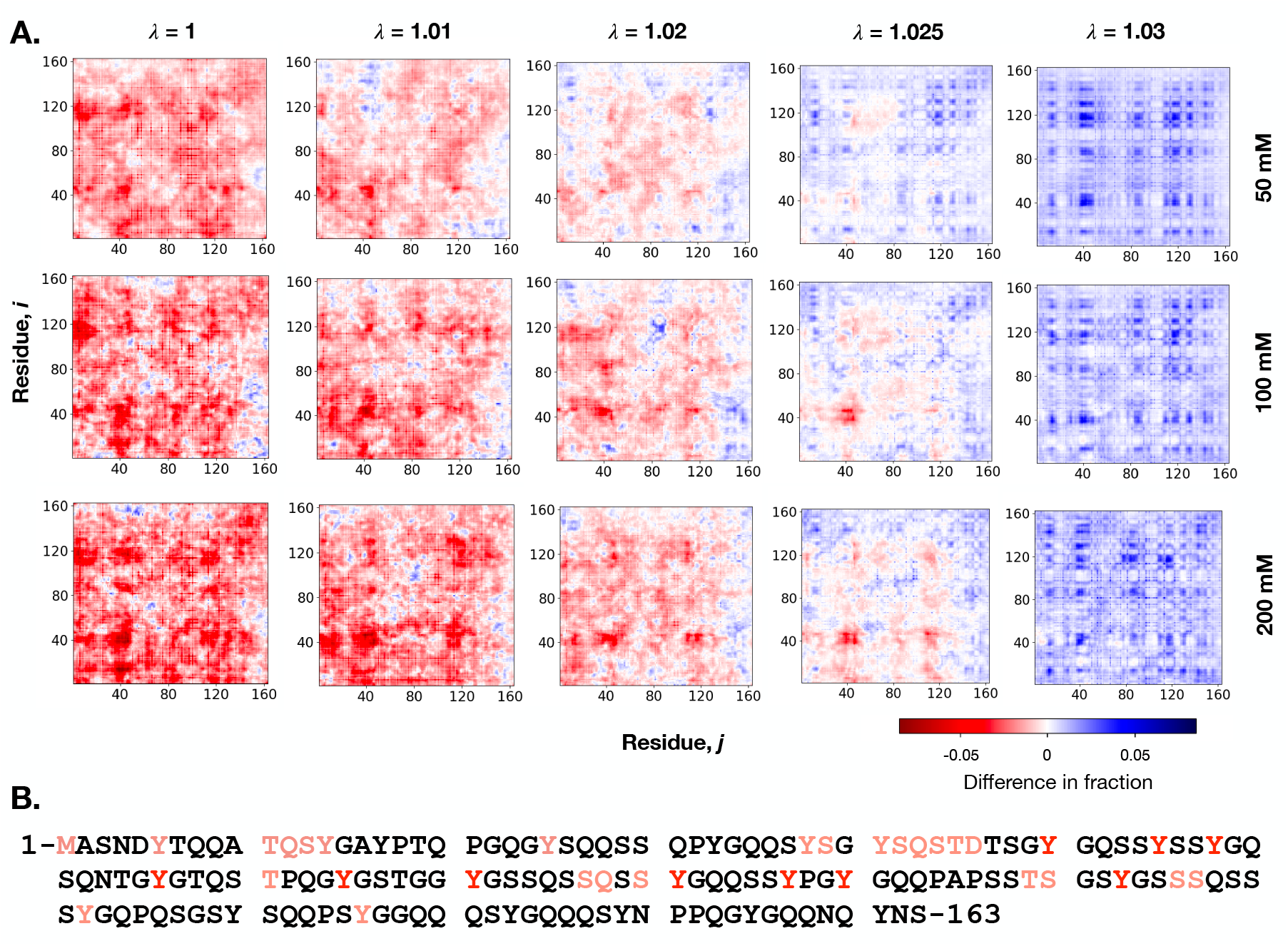
Summary of the effect of salt concentration on the inter-chain contacts. A. The difference in the fraction of the inter-chain residual contacts at various salt concentrations with respect to 0 mM salt. The color axis corresponds to (contact fraction|_X mM_ - contact fraction|_0 mM_) where X is 50, 100, or 200 mM salt concentrations. B. FUS LC amino acid sequence. Amino acids with a high propensity to form contacts at 0 mM salt concentration for *λ*=1 (fraction > 0.1 in the top left corner of Figure 5) are shown in red font. The ones shown in faded red font are the ones that have fraction > 0.1 at 0 mM salt concentration but lose the strength to have fraction < 0.1 as the salt concentration increases (when *λ* < 1.03, as the salt makes all contacts stronger at *λ*=1.03 for all salt concentrations.

Figure 6A also shows the effect of salt concentration on inter-chain contact formation (from top to bottom for each *λ*). We found that the contact formation profile has substantially changed upon the addition of salt. At *λ*=1, where we found the largest concentration for the dense phase (Figure 4), the increase in salt concentration systematically reduces the contact fraction (Figure 6A, leftmost column). This reduction is more prominent in the fraction of cluster-type contacts (for example, between residues 35 to 50). Interestingly, however, contact clusters reappear in the presence of as the *λ* increases (Figure 5). FUS LC domain has characteristics of prion-like proteins, including being predominantly composed of uncharged amino acids.^49^ FUS LC domain has only 2 charged residues out of 163.These charged residues are located at positions 5 and 46, where we observe most of the reduction in the fraction of cluster-type contacts.

As the *λ* increases, however, the effect of salt reverses its trend, that is, the addition of salt makes the contact fraction weaker until *λ*=1.02-1.025, the change in contact fraction becomes more balanced at *λ*=1.02-1.025, but at *λ*=1.03, all contacts become stronger with the addition of salt. Previously, a decreased solubility of FUS LC upon an increase in salt concentration was reported by Fawzi and coworkers.^36^ The authors have reasoned that the salting-out of the hydrophobic residues (predominantly tyrosines) could be responsible for the decreased solubility. Our finding of the increase in protein-protein contacts at *λ*=1.03 is consistent with this salting-out observation. Although we do not directly measure solubility, the salting-out effect means an effective increase in protein-protein interactions, which is consistent with the increase in amino acid contact fractions as we observe here. We note that the *λ*=1.03 is the strength of protein-water interactions that gives an agreement in the transfer free energy with the experimental values for the systems with salt. Therefore, it is reasonable that we are observing the experimentally predicted effects at this specific *λ*, *λ*=1.03. This observation provides another layer of validation to our modeling approach.

For the smaller *λ* values, protein-protein interactions dominate the protein-water interactions. As opposed to the net increase in amino acid contacts (i.e., salting-out) observed at higher *λ* (*λ*=1.03), we observed an effective reduction in amino acid contact fraction at lower *λ* (Figure 6), which is rather consistent with electrostatic screening. We note that even though protein-protein contact fractions become lower upon the addition of salt (compared to 0mM salt) at low *λ*, this reduction neither causes a dissociation of condensates (Figure 1, blue data, left column) nor a significant change in the protein concentration in the dense phase (Table 1). There are, however, significant changes to the condensate morphology. Increased salt concentration at low *λ* yields percolated nonspherical protein condensates instead of spherical droplets (Figure 2). The shape of the condensed phase is governed by surface tension. Surface tension (or interfacial tension) arises from the balance between intermolecular forces:^50^ If the cohesive forces are stronger than the adhesive forces, protein molecules will collapse (or coalesce) into a condensed phase due to the inward component of the net force. Moreover, this condensed phase will also tend to shrink its interface due to the tangential component of the net force. Interfacial tension causes the biomolecular condensates to minimize their surface area with the dilute phase, driving them towards a spherical shape (since a sphere provides the minimum surface for a given volume). Accordingly, we argue that the deviations from spherical shapes at elevated salt concentrations (at low *λ*, especially *λ*=1) may be explained by lower surface tension that arises from effective protein-protein interactions being reduced with the presence of salt.

## Conclusions

In this work, we studied the condensate formation of the FUS LC domain at various salt concentrations using a MARTINI 3.0 force field by systematically scaling the interaction strength of protein-water beads, introducing a parameter, *λ*. *λ* is a factor that we multiply the e of the Lennard-Jones interaction between protein-water beads, that is the larger the *λ* is, the stronger the protein-water interactions are, and *λ*=1 is the base strength of protein-water interactions. We systematically increased the *λ* in order to reproduce the condensate properties, specifically the excess free energy of transferring protein chains from the dilute phase to the condensed phase. While we do not observe the formation of a dilute phase at low protein-water interactions, that is, all protein chains coalesce into one dense phase, when the protein-water interactions are unmodified, or when the *λ* < 1.02, as we scaled *λ* up, we started observing a phase equilibrium. We presented the results from each scaling at various salt concentrations. The optimal *λ* is slightly different in the presence or in the absence of the salt. But within the concentration range of the salt that we tested for the FUS LC domain, the optimal *λ* does not change at different salt concentrations; it only changes based on whether we have salt in the system or not.

We reported a range of *λ* values where we were able to reproduce experimentally observed transfer free energy. Although we were able to reproduce the transfer free energy, we found that the protein concentration in the dense phase was still higher than the experimentally detected concentrations. We also found that the presence of salt concentration affects the dense phase morphology, especially when protein-water interactions are weaker (low *λ*). While a more extensive exploration of the effect of salt concentration on condensate morphology is a part of future work, here, we showed that the individual amino acid contacts are highly sensitive to salt concentration, which might be related to nonspherical morphology. Salt has more predominant screening effects at low *λ* whereas the opposite salting-out effects become more predominant as *λ* increases. While the screening effects do not cause dissociation of the condensates, we argued that the screening may be responsible for condensates losing their spherical shape by reducing the surface tension. At high *λ*, we did not observe any obvious effects of salt on morphology, where the salting-out effects are more predominant. In summary, our findings show that there is a balance between electrostatic screening effects and salting-out effects and this balance is highly sensitive to the strength of protein-water interactions (*λ*).

## Acknowledgement

This work is partly supported by funding from the Cancer Prevention and Research Institute of Texas (CPRIT) award RR220008. GHZ is deeply grateful to Prof. Pablo Debenedetti of Princeton University for his support (including but not limited to financial and computational) in the early course of this work. GHZ also thanks Dr. Hasan Zerze of the University of Houston for the assistance with OVITO analyses and contact map calculations. The simulations presented in this work are performed partly on computational resources managed and supported by Princeton Research Computing (a consortium of groups including the Princeton Institute for Computational Science and Engineering [PICSciE] and the Office of Information Technology’s High Performance Computing Center and Visualization Laboratory at Princeton University) and partly on the computational resources provided by the Hewlett-Packard Enterprise Data Science Institute in the University of Houston.

## Supporting Information Available

3 Supporting Figures, 1 Supporting Table, and Supporting Text are available.

## Graphical TOC Entry

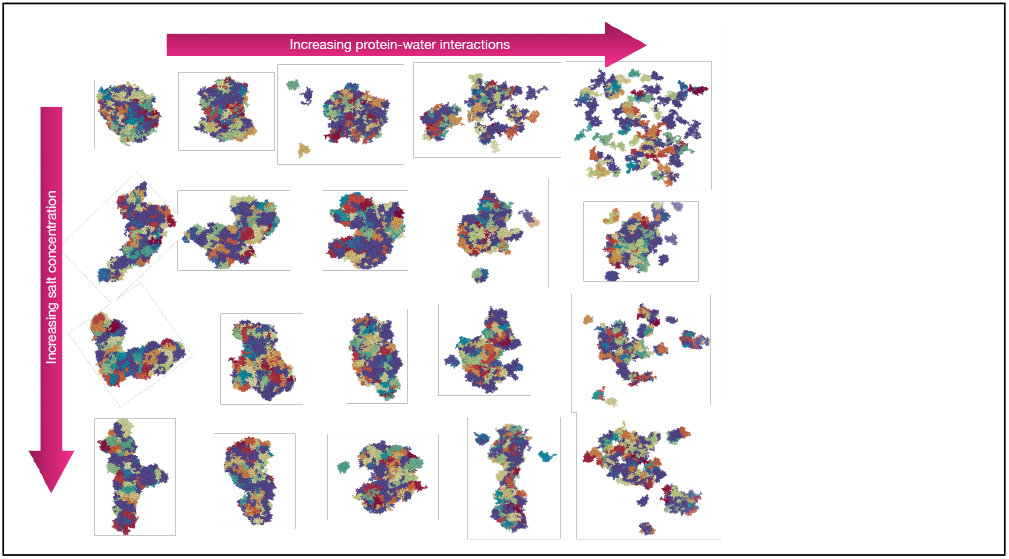

## Notes

### Competing Interest Statement

The authors have declared no competing interest.

