## Supplementary Information for "Optimizing the Martini 3 force field reveals the effects of the intricate balance between protein-water interaction strength and salt concentration on biomolecular condensate formation"

Gül H. Zerze*

*William A. Brookshire Department of Chemical Biomolecular Engineering, University of Houston, Houston, TX, 77204, USA*

#### Supporting Text

##### FUS LC sequence

1-MASNDYTQQA TQSYGAYPTQ PGQGYSQQSS QPYGQQSYSG YSQSTDTSGY GQSSYSSYGQ SQNTGYGTQS TPQGYGSTGG YGSSQSSQSS YGQQSSYPGY GQQPAPSSTS GSYGSSSQSS SYGQPQSGSY SQQPSYGGQQ QSYGQQQSYN PPQGYGQQNQ YNS-163

#### Supporting Figures

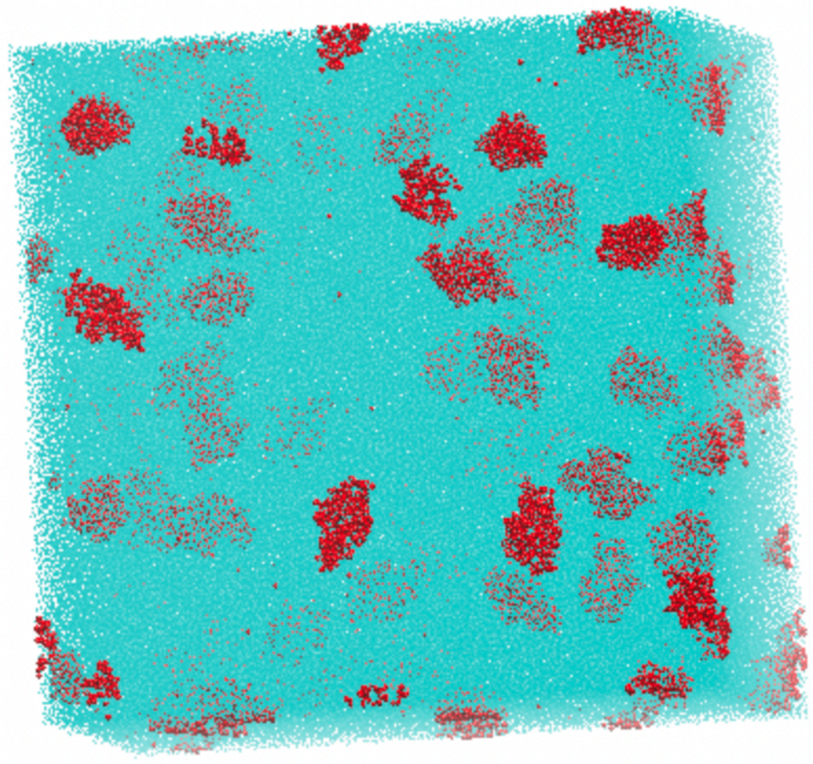

Figure S1: A snapshot of an example monodispersed initial configuration.

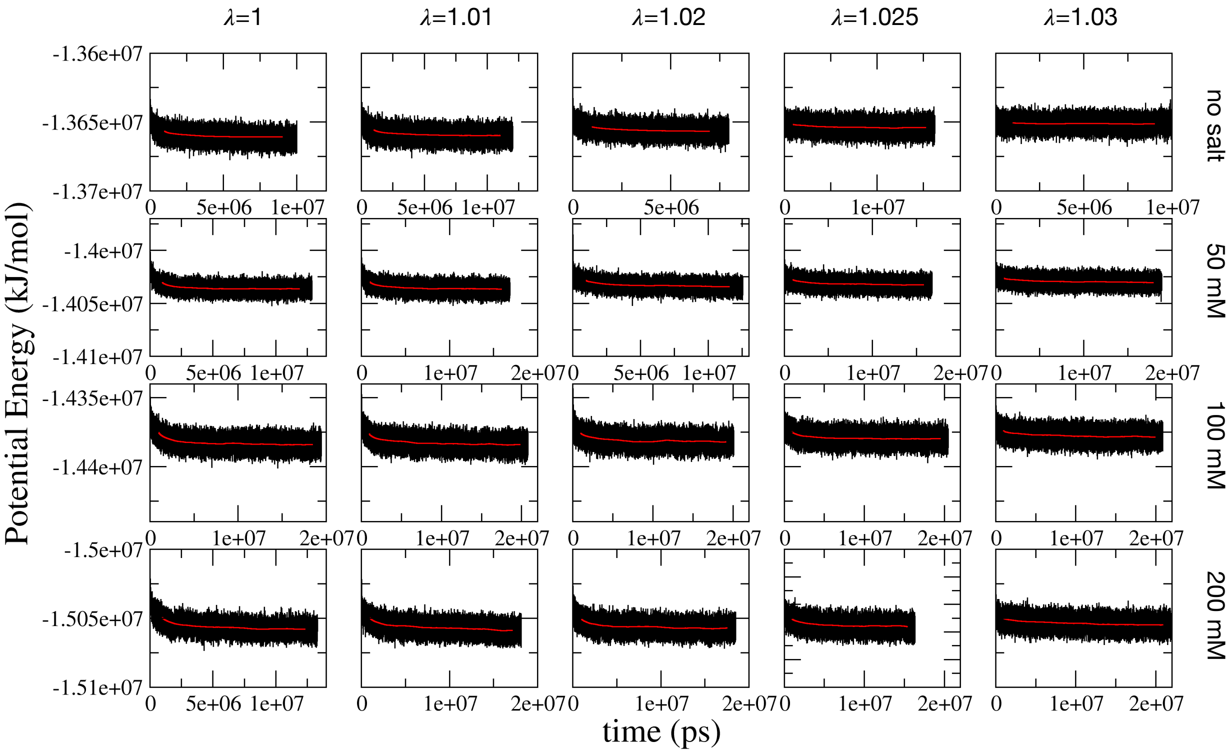


Figure S2: Total potential energy of systems as a function of time (black) and its running average (red).

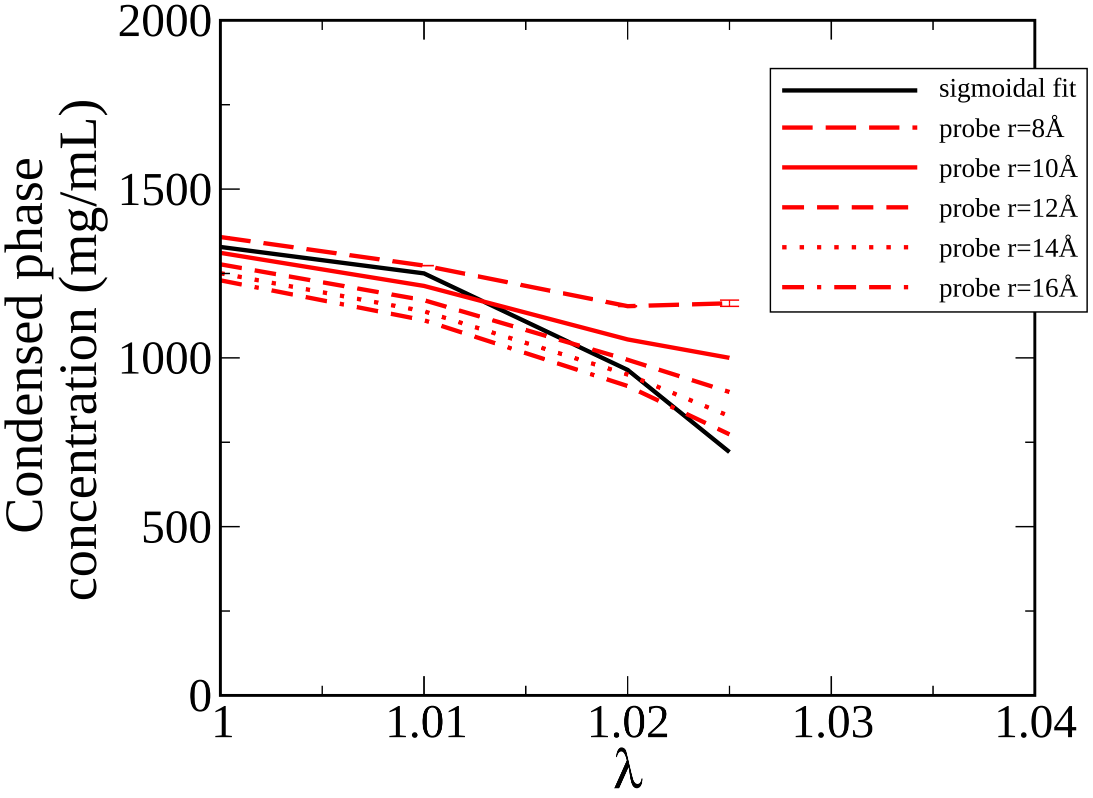


Figure S3: Protein concentration in the condensed phase analyzed by two different techniques (sigmoidal fit: black, surface reconstruction: red). For the surface reconstruction technique, different probe radii tests are shown in broken lines.

### Supporting Tables

Table S1: Equilibration time periods for all systems in units of ps.

| | $\lambda = 1$ | $\lambda = 1.01$ | $\lambda = 1.02$ | $\lambda = 1.025$ | $\lambda = 1.03$ |
|---|---|---|---|---|---|
| **no salt** | 7,000,000 | 6,000,000 | 7,000,000 | 8,000,000 | 5,000,000 |
| **50 mM salt** | 3,000,000 | 7,000,000 | 9,000,000 | 10,000,000 | 10,000,000 |
| **100 mM salt** | 6,000,000 | 6,000,000 | 6,000,000 | 10,000,000 | 10,000,000 |
| **200 mM salt** | 7,000,000 | 10,000,000 | 9,000,000 | 6,000,000 | 15,000,000 |
